# Association of Facial Aging with DNA Methylation and Epigenetic Age Predictions

**DOI:** 10.1101/367326

**Authors:** Riccardo E Marioni, Daniel W Belsky, Ian J Deary, Wolfgang Wagner

## Abstract

Evaluation of biological age, as opposed to chronological age, is of high relevance for interventions to increase healthy aging. Highly reproducible age-associated DNA methylation (DNAm) changes can be integrated into algorithms for epigenetic age predictions. These predictors have mostly been trained to correlate with chronological age, but they are also indicative for biological aging. For example accelerated epigenetic age of blood is associated with higher risk of all-cause mortality in later life. The perceived age of facial images (face-age) is also associated with all-cause mortality and other aging-associated traits. In this study, we therefore tested the hypothesis that an epigenetic predictor for biological age might be trained on face-age as surrogate for biological age, rather than on chronological age. Our data demonstrate that facial aging and DNAm changes in blood provide two independent measures for biological aging.

## Main Text

We analysed data from the Lothian Birth Cohort 1921 (LBC1921), a longitudinal study of aging in a 1921 birth cohort followed up at five assessment waves between ages 79 and 92 years [1]. DNA methylation profiles were analysed in whole blood collected at age 79.1 years using Illumina HumanMethylation450BeadChips as previously described [2]. Perceived facial age (face-age) was assessed from neutral expression facial photographs taken at age 83.3 years (blood samples were not collected at this time point)[3]. Briefly, 12 university students (6 male, 6 female) estimated the participants ages based on high resolution photographs, taken under the same lighting conditions, at the same distance, using the same camera. The images were presented one at a time on a high-quality cathode ray tube computer monitor. Face-age acceleration was calculated as the (linear regression) residuals of face-age regressed on chronological age. Mean estimated face-age was 74.2 years (SD 3.9, range 63.5–85.3). Overall, DNAm measurements and face-age assessments were available for 235 individuals (43% female; 6% current smokers; 49% ever smokers).

Perceived face-age has been linked to mortality risk [4] and other ageing-associated traits [5]. The relationship between older face-age and increased mortality risk was also evident in a previous data release of LBC1921[3], and again here using updated survival information (HR 1.39 [1.19, 1.63] per SD increase in face-age; **Figure 1A**). People with older face-age also show signs of accelerated biological aging as measured from physiology-based indices (Pearson r~0.2) [6]. Evidence for association of older face-age with epigenetic aging is more sparse; in midlife adults of the Dunedin study there was a small effect-size association with one epigenetic clock but no association with two others [6]. We therefore first tested associations between face-age and the three epigenetic age predictors proposed by Horvath (based on 353 CG dinucleotides - CpGs) [7], and two signatures that were trained on blood samples by Hannum et al. (71 CpGs)[8] and Weidner et al. (99 CpGs) [9]. In the LBC1921 cohort, consisting of older adults, accelerated epigenetic aging (residual of epigenetic age regressed on chronological age) was not associated with higher face-age (r_Horvath:face-age_=0.06, P=0.35; r_Hannum:face-age_=0.01, P=0.93; and r_Weidner:face-age_=0.01, P=0.82).

**Figure 1:**
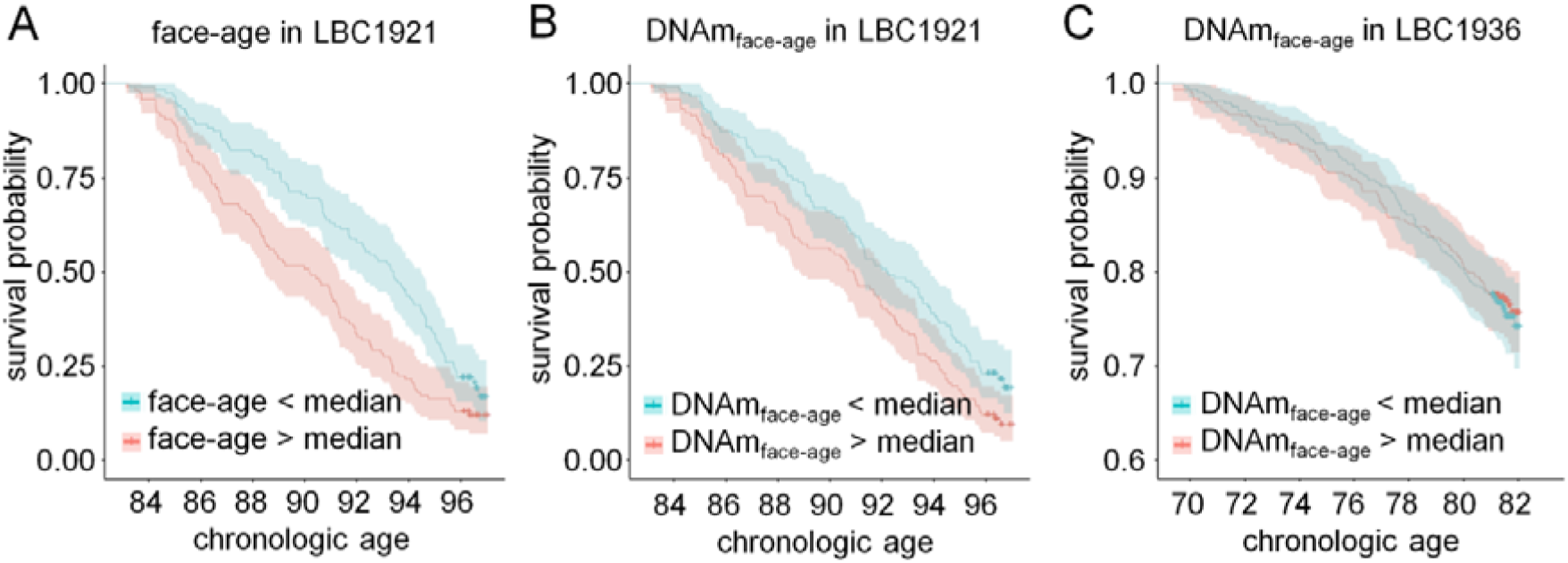
Perceived facial aging is associated with all-cause mortality, but not with DNA methylation signatures trained on perceived aging. **(A)** Kaplan-Meier Plots depict survival rates of LBC1921 participants stratified by the median perceived age in facial images (face-age). **(B)** Alternatively the participants were stratified by mean age-predictions based on an algorithm of 32 CpGs that was trained on face-age of the LBC1921 (DNAm_face-age_). **(C)** The results with this algorithm did not replicate in the independent LBC1936 cohort.

We then tested the associations between face-age and DNAm of 307,745 individual CpGs (Table S1). The maximum absolute Spearman correlation of face-age with DNAm was r=0.29 (P=5.1×10^−6^) for cg18402261. To take potential confounding effects into account we then performed epigenome-wide association study (EWAS) analysis with three nested regression models: the base model (M1; **Table S2**) included covariates for age, sex, and technical factors (plate, array, position, hybridisation date); additional models added covariates for smoking history (M2; **Table S3**) and measured white-blood-cell counts (neutrophils, lymphocytes, monocytes, eosinophils, and basophils; M3; **Table S4**). The lead CpG from the most conservative model (M3) was cg00871706 (P=1.8×10^−6^, chr7:138,666,647, *KIAA1549*). Pathway analysis of the top 100 CpGs from this most conservatively-modelled EWAS identified no evidence of functional enrichment (Bonferroni P>0.05). Taken together, the perceived age based on facial photographs revealed no genome-wide significant associations with DNAm at specific CpG sites or with gene sets linked to specific biological pathways.

Finally, we tested if a combination of the top 100 CpGs might provide information about biological aging. We conducted LASSO regression analysis to fit data from the top 100 CpGs in the M3 EWAS to facial age. The resulting algorithm included 32 CpGs (**Table S5**). In the LBC1921 training sample (n=235, n_deaths_=198), the epigenetic predictor of facial age was correlated with measured facial age (r=0.66) and predicted increased risk for mortality (HR 1.31 [1.12, 1.53] per SD increase in face-age; **Figure 1B**). The association with mortality risk was in the same direction but had a much smaller effect-size and was not statistically significant in the out-of-sample analysis in the independent Lothian Birth Cohort 1936 (n=920, n_deaths_=215; HR 1.07 [0.94, 1.23], p=0.32; **Figure 1C**).

This is the first study on epigenome-wide association of DNAm with perceived facial age. Face-age was clearly associated with all-cause mortality in the 235 LBC1921 individuals, as previously reported for an older iteration of the data [3]. However, our results did not support the hypothesis that an epigenetic measure of biological age could be derived from analysis of facial aging. Efforts to train epigenetic predictors of biological aging on surrogate biological aging measures may require larger sample numbers. On the other hand, precise epigenetic aging signatures have been trained for chronological age on smaller sample sets [9]. A second limitation is that face-age and DNAm were measured in skin and blood, respectively. It is well known that age-associated modifications occur in both tissues [7], but the pace of biological aging may be independent in different tissues. DNAm analysis in skin should therefore be a priority in future face-age analyses; however, such datasets are not yet available. Our results indicate that face-age and epigenetic aging signatures of blood provide independent and complementary measures for biological age.

## Materials and Methods

### The Lothian Birth Cohorts

The Lothian Birth Cohort 1921 (LBC1921) is a longitudinal study of aging [1]. All participants were born in 1921 and have been followed up every few years from ages 79 to 92 years, yielding a maximum of 5 waves of data per participant. At each wave, cognitive, personality, health and disease data were collected.

The Lothian Birth Cohort of 1936 (LBC1936) includes participants that were born in 1936 and have been followed up every three years from ages 70 to 82 years [1]. The most recent wave of data collection at age 82 years is currently ongoing. As with the LBC1921 study, cognitive, personality, health and disease data were collected at each wave.

Survival data are routinely collected in LBC via data linkage to the Scottish National Health Service Central Register. Data for this study were correct as of January 2018.

### DNA methylation

DNA methylation data were assessed in whole blood samples from the LBC studies using the Illumina HumanMethylation450 BeadChip (Illumina Inc., San Diego, CA). Data from the first wave of data collection in both LBC studies were considered for the current analysis: LBC1921 participants were a mean age of 79.1 years (SD 0.55 years); LBC1936 participants were a mean age of 69.5 years (SD 0.83 years). Background correction was performed and quality control was used to remove probes with a low detection rate (P>0.01 for >5% of samples), low quality (manual inspection), low call rate (P<0.01 for <95% of probes), and samples with a poor match between genotypes and SNP control probes, with incorrectly predicted sex. This left a dataset of 450,727 CpGs, which was further filtered to exclude X and Y, cross-reactive, non-cg, and SNP-in-probe-sequence CpGs, The final EWAS dataset therefore consisted of 307,645 CpGs.

### Face-age and DNAm age acceleration correlations

Pearson correlations were computed between the age acceleration measures (face-age from age 83 years and DNAm age from age 79 years).

### Epigenome-wide association studies

Three linear regression models were considered for the EWAS as described in the text. The DNAm values at individual CpGs were the dependent variables and face-age was the dependent variable of interest. A Bonferroni correction was applied to account for multiple testing (P<0.05/307,745=1.6×10^−7^).

### Functional enrichment

The top 100 CpGs from the most conservative EWAS model (M3) were examined for functional enrichment (closest gene to the nearest transcription start site was taken as the input) based on a PANTHER Gene Ontology analysis with Bonferroni P<0.05 set as the significance threshold.

### LASSO regression

A parsimonious predictor of face-age was then generated via least absolute shrinkage and selection operator (LASSO) regression on the 100 CpGs most associated with face-age from the M3 EWAS using the ‘glmnet’ package in R. Mean imputation was used for any missing CpG values (n=3). 10-fold cross-validation was applied with the mixing parameter (alpha) set to 1 (LASSO penalty). Coefficients were extracted for the model with the minimum mean cross-validated error estimate.

Age- and sex-adjusted Cox proportional hazards models with DNAm-based face-age as the predictor and time-to-event (death or censoring) as the outcome were performed in the training and test datasets (LBC1921 and LBC1936, respectively). Analyses were conducted in R using the ‘survival’ package.

## Declarations

### Ethics approval and consent to participate

Following written informed consent, peripheral whole blood was collected for DNA extraction in LBC1921. Ethics permission for the LBC1921 was obtained from the Lothian Research Ethics Committee (LREC/1998/4/183 and LREC/2003/7/23). LBC1936 ethics permission was obtained from the Multi-Centre Research Ethics Committee for Scotland (MREC/01/0/56) and the Lothian Research Ethics Committee (LREC/2003/2/29).

### Availability of data and material

LBC data are available upon request: https://www.lothianbirthcohort.ed.ac.uk/content/collaboration

### Author contributions

REM, DWB, and WW designed the study and analysed the data. All authors contributed to the writing of the manuscript.

## Acknowledgements

We thank Mrs Alison Pattie for her help in collecting data from, and assisting in the administration of, the LBC1921 cohort. We thank other Lothian Birth Cohort team members for assistance with data collection and collation.

